# *Cis* and *trans*-acting variants contribute to survivorship in a naïve *Drosophila melanogaster* population exposed to ryanoid insecticides

**DOI:** 10.1101/502161

**Authors:** Llewellyn Green, Paul Battlay, Alexandre Fournier-Level, Robert T. Good, Charles Robin

## Abstract

Insecticide resistance is a paradigm of microevolution and insecticides are responsible for the strongest cases of recent selection in the genome of *Drosophila melanogaster*. Here we use a naïve population and a novel insecticide class to examine the *ab initio* genetic architecture of a potential selective response. Genome wide association studies of chlorantraniliprole susceptibility reveal variation in a gene of major effect, *Stretchin Myosin light chain kinase* (*Strn-Mlck*), which we validate with linkage mapping and transgenic manipulation of gene expression. We propose that allelic variation in *Strn-Mlck* alters sensitivity to the calcium depletion attributable to chlorantraniliprole’s mode of action. Genome-wide association studies also reveal a network of genes involved in neuromuscular biology. In contrast, phenotype to transcriptome associations identify differences in constitutive levels of multiple transcripts regulated by cnc, the homologue of mammalian Nrf2. This suggests that genetic variation acts in *trans* to regulate multiple metabolic enzymes in this pathway. The most outstanding association is with the transcription level of *Cyp12d1* which is also affected in *cis* by copy number variation. Transgenic overexpression of *Cyp12d1* reduces susceptibility to both chlorantraniliprole and the closely related insecticide cyantraniliprole. This systems genetics study reveals multiple allelic variants segregating at intermediate frequency in a population that is completely naïve to this new insecticide chemistry and it adumbrates a selective response among natural populations to these chemicals.

**Significance:** Around the world insecticides are being deregistered and banned, as their environmental costs are deemed too great or their efficacy against pest insects is reduced through the evolution of insecticide resistance. With the introduction of replacement insecticides comes the responsibility to assess the way new insecticides perturb various levels of biological systems; from insect physiology to ecosystems. We used a systems genetics approach to identify genetic variants affecting survivorship of *Drosophila melanogaster* exposed to chlorantraniliprole. The study population was completely naïve to this insecticide chemistry and yet we find associations with variants in neuromuscular genes and co-regulated detoxification genes. We predict that these variants will increase in populations of this ‘sentinel species’ as these insecticides are applied in the environment.

## Introduction

An elaboration of the adage of Paracelsus (1493-1541) that ‘the dose makes the poison’ is that there is a dose range of insecticides that kills some but not all insects in a population. By examining the genetic variation that contributes to survivorship on such discriminating doses, we can take a genetics approach to address a diverse set of questions relating to insecticide biology. Which genes have variants that affect survivorship, and how do they combine to provide the genetic architecture underpinning the trait? Do they provide insights into the mode of action of new insecticides? Do they suggest likely mechanisms by which insecticide resistance will arise? And what else do they tell us about past and future evolutionary responses to insecticides in pest and non-target species?

Chlorantraniliprole (Rynaxapyr), is the first of the anthranilic diamides, a new class of insecticides. Unlike earlier insecticides that predominantly target neurotransmission (1, 2), the anthranilic diamides are designed to target the ryanodine receptor, which is primarily involved in calcium homeostasis and muscle contraction. Disruption of ryanodine receptor activity provides a rapid incapacitation of the pest, leading to feeding cessation, lethargy, paralysis, and death (3-5). Therefore, both the mode of action and the chemistry suggest that cross-resistance with older insecticides is unlikely.

Chlorantraniliprole was first sold in the Philippines in 2007, and worldwide soon after (6). Within years of introduction, resistance cases were reported in the diamondback moth *Plutella xylostella* (7-9), and the tomato leafminer *Tuta absoluta* (10). While some of these cases can be attributed to mutations in the ryanodine receptor, the primary molecular target of these insecticides, there are others that suggest that resistance to this new insecticide class can arise through other means (11).

While *Drosophila melanogaster* is not a pest or a direct target of chlorantraniliprole applications, it is an organism of interest for two reasons. Firstly, *D. melanogaster* has long served as a model for insecticide resistance (12) and its status as a model organism more generally means that there are a wide variety of tools available to characterize genetic traits (13). Secondly, selective sweep analyses show that insecticides (particularly the organophosphates) have been major selective agents on *D. melanogaster* populations (14-17). These findings support the proposition that *D. melanogaster* can be used as a sentinel species for environmental pollutants, particularly insecticides (18).

Like pest insects, *D. melanogaster* evolves insecticide resistance chiefly through target molecule insensitivity or detoxification enzyme adaptation, although other resistance mechanisms have been characterized (19). Resistance mutations in the target site genes typically diminish insecticide binding (e.g. amino acid substitutions in *Ace* confer organophosphate and carbamate resistance; 20). Resistance mutations affecting detoxification enzymes can alter the protein sequence (eg. *Cyp6w1;* 21) but more generally increase transcriptional output through copy number variation or *cis*-regulatory changes in the promoters of the resistance genes (eg. up-regulation of P450 enzymes such as *Cyp6g1*; 22, 23). However, master regulatory genes that control, in *trans*, detoxification pathways have been speculated to be involved in resistance in various pest species (24) as well as in *D. melanogaster* (25) but have so far not been well characterized.

A recent addition to the *Drosophila melanogaster* toolkit is the Drosophila Genetic Reference Panel (DGRP; 26), which comprises 205 inbred lines derived from a single North American population. Each line in the DGRP has been bred for homozygosity and its genome sequenced, creating a ‘living library’, designed for genome wide association studies that will associate genetic variants with phenotypes. The DGRP has been phenotyped for an extensive number of traits (27) including various insecticide phenotypes (15-17, 21, 28, 29) and ‘intermediate phenotypes’ such as transcript abundance that enables eQTL to be mapped (30). Thus, the DGRP is becoming an important model for systems genetics of insects (31, 32), which, as we demonstrate here, enables deeper characterization of the genetic and regulatory mechanisms underpinning traits.

The DGRP lines were established from a collection of flies from the Farmers Market in Raleigh, North Carolina (USA) in 2003 (26), prior to the introduction of chlorantraniliprole into the market in 2007. Thus, they are naïve with respect to the completely novel class of chemistry of the group 28 insecticides. This provides a rare opportunity to examine the *‘ab initio’* state of a potentially adaptive trait at unprecedented genetic resolution.

Here, we investigate the genetic architecture of chlorantraniliprole susceptibility across multiple doses. We interrogate associations between chlorantraniliprole phenotypes and both genomic and transcriptomic variation. We find that an allele of large effect already segregates within this population, which is detectable only at higher levels of exposure. In addition, we show that phenotype to transcriptome associations reveal a completely different set of candidate genes, linked by a common *trans*-regulatory pathway, which in cases of insecticide biology has rarely been demonstrated at a molecular level.

## Results

### Phenotype to genome associations (GWAS)

154 DGRP lines were scored for larval survivorship on six doses of chlorantraniliprole (Fig. 1A). The broad-sense heritability (H^2^) of the six doses ranged from 0.73–0.85, indicating a strong genetic component to chlorantraniliprole survivorship across the DGRP. Genome-wide association studies (GWAS) on each of the doses identified 42–335 variants below the arbitrary genome-wide significance threshold (*p*<1×10^-5^). The number of variants passing this threshold increased with dose; however this relationship failed to account for linkage disequilibrium. Therefore, we examined the explanatory power of phenotype-associated DGRP variants for each chlorantraniliprole dose using a multivariate genomic prediction model and found that more genes are required to explain the genetic architecture of the lowest dose (0.5μg/mL). For example, if the top 50 most-associated variants of the 0.5ug/mL dose are considered they explain about the same amount of the phenotypic variation (R=0.43) as the top five variants of the 5ug/mL dose (Fig. S1).

**Figure 1.**
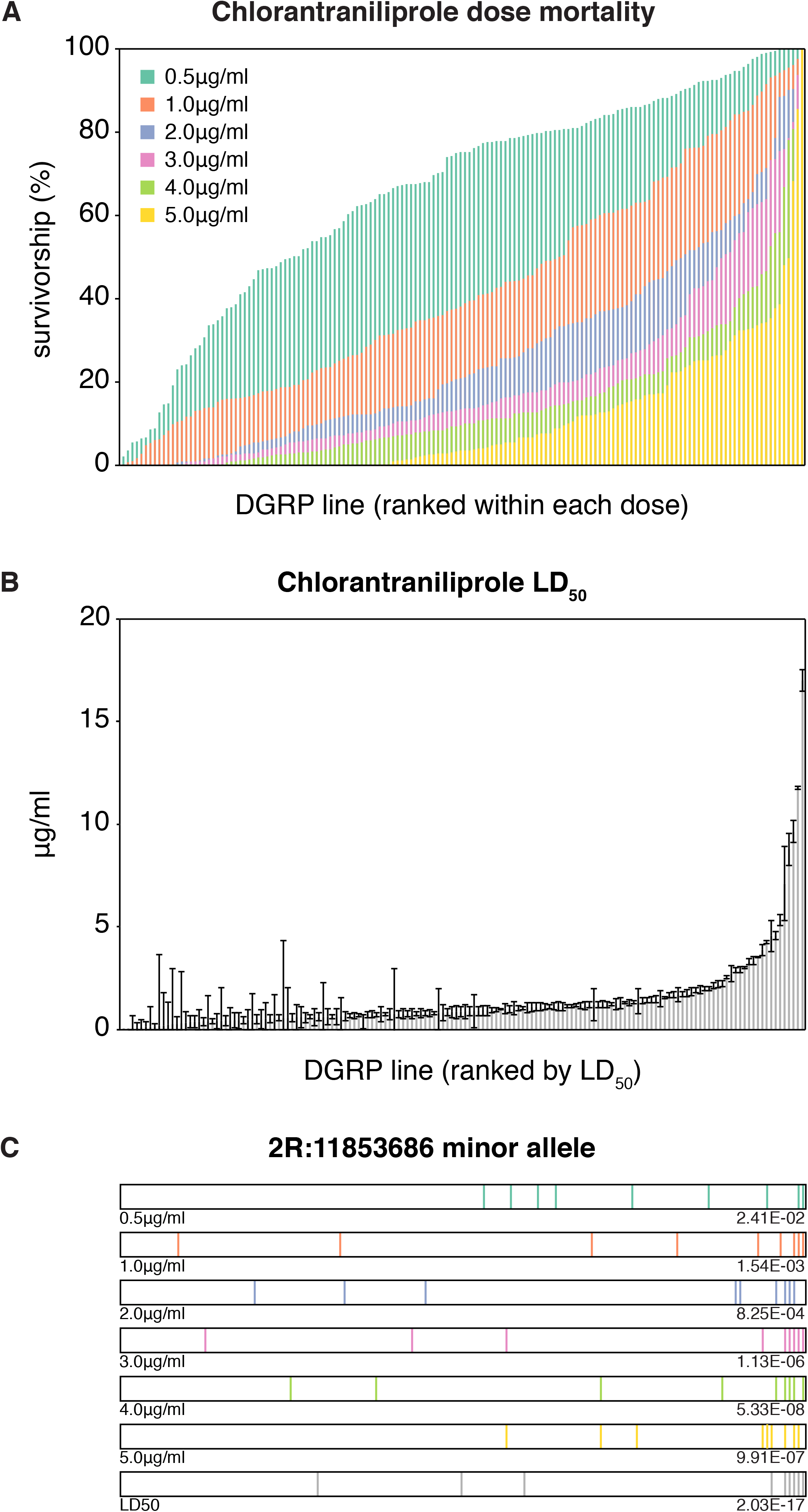
Chlorantraniliprole phenotypes of 154 DGRP lines. (*A*) The survivorship of each line on the different doses (coloured). The lines are ordered by LD_50_ survivorship. (*B*) LD_50_, calculated from a minimum of six Abbott-corrected dose survivorship phenotypes. (*C*) The distribution and minimum p-value of the top candidate (*Strn-Mlck;* 2R:11853686) from the LD_50_ GWAS across doses.

These data were supplemented with screening at additional doses for some DGRP lines to estimate the dose of chlorantraniliprole required to kill 50% of individuals in each line (LD_50_). The mean LD_50_ was equal to 1.4μg/mL (SD=2.07μg/mL) while the maximum LD_50_ was 17ug/mL (Fig. 1B). A GWAS of the LD_50_ phenotype identified 931 associated variants below the arbitrary genome-wide significance threshold (*p*<1×10^-5^; Fig. 2A). Ninety-six of these remained significant after a Bonferroni correction for multiple testing (2.65×10^-8^; Supplementary file). The strongest association was to a single nucleotide polymorphism (SNP) in an intron of *Stretchin myosin light chain kinase* (*Strn-Mlck*; 2R:11853686; *p*=2.03×10^-17^; Fig. 1C, Fig. 2), and variants annotated to this gene accounted for 15 of the 96 Bonferroni-significant GWAS variants. Change in genetic architecture with respect to dose is also illustrated when 2R:11853686 is examined at other dose phenotypes where it is not associated with a detectable increase in survivorship (Fig. 1C).

**Figure 2.**
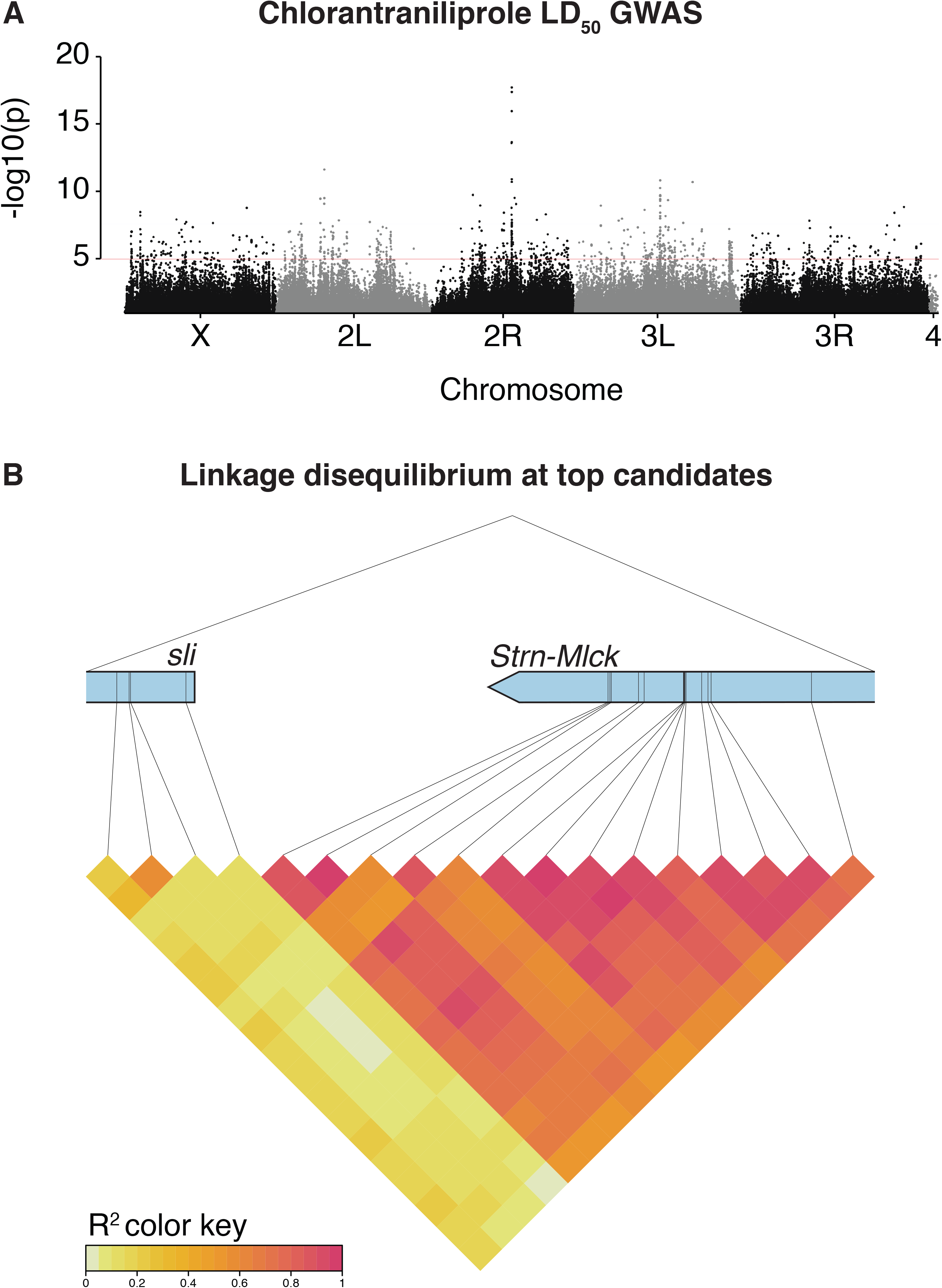
(*A*) Top associations with genomic variants. Manhattan plot (-log_10_(*p*) against genomic position) of DGRP chlorantraniliprole LD_50_ associations. *Strn-Mlck* is a standout candidate, with a minimum *p*-value of 2.03×10^-17^. (B) Linkage Disequilibrium at LD_50_ GWAS top candidates *sli* and *Strn-Mlck*.

To account for the fact that the strong effect of *Strn-Mlck* variation may be influencing other GWAS associations, we fitted the effect of *Strn-Mlck* to the chlorantraniliprole LD_50_ data and ran a GWAS on the corrected phenotype. The gene with the most highly associated variants after *Strn-Mlck* in the original LD_50_ GWAS, *sli*, was maintained in the *Strn-Mlck-corrected* GWAS. This demonstrates that despite its proximity to *Strn-Mlck*, associations with variants in *sli* are not artefacts of linkage disequilibrium with variants in *Strn-Mlck* (Fig. 2B, Fig. S2).

*sli* is like other genes harbouring highly associated variants in that it is involved in axon guidance, sarcomere organization, intracellular signalling and regulation of cell growth. The R spider software (33) was used to identify networks of genes in the Kyoto Encyclopedia of Genes and Genomes (KEGG) or Reactome pathways that were enriched for variants associated with chlorantraniprole phenotypes. The top 100 variants from LD_50_ genomic prediction both before and after correcting for the *Strn-Mlck* association were considered. The uncorrected LD_50_ network contained 12 genes, five with GWAS associations, out of a total of 54 recognized by R spider (Monte Carlo simulation p=0.01; Fig. S3); the Strn-Mlck-corrected LD_50_ network contained 10 genes, five with GWAS associations, out of a total of 53 genes with annotated function, of which 7 could be mapped to a reference network previously described in KEGG (p=0.005; Fig. S3). Three GWAS-associated genes appeared in both networks: *sli, robo3* and *norpA*. Although not connected to either network based on Reactome or KEGG databases, *Strn-Mlck* can be linked to the corrected LD_50_ network, as it has been shown to be one of the serine/threonine kinases that phosphorylates the transcription factor *foxo* (34).

### Phenotype to transcriptome associations

Transcript-level variation from the DGRP transcriptome dataset (mean for males and females; 30) was associated with chlorantraniliprole LD_50_ for 9 genes below Bonferroni significance (GLM *p*<2.76×10^-6^; Fig. 3). These top candidates are enriched for members of genetically correlated transcription modules (30) for both males (module 79; 7 transcripts) and females (module 21; 6 transcripts). Furthermore, eight of the nine Bonferroni-significant transcripts have also been shown to be induced through ectopic expression of the transcription factor *Cap ‘n’ collar* (*cnc*) and following exposure to phenobarbital (Fig. 3; 35). One top candidate, *Cyp12d1*, exhibits copy number variation (CNV) within the DGRP such that two copies (*Cyp12d1-p* and *Cyp12d1-d*) are observed in 24% of lines. The DGRP was explicitly genotyped for duplication of *Cyp12d1* and the duplication was found to be correlated with *Cyp12d1* transcription levels (*Cyp12d1-p:* male r^2^=0.12, female r^2^=0.14; *Cyp12d1-d:* male r^2^=0.38, female r^2^=0.27). Thus, two distinct contributions to *Cyp12d1* transcript level can be identified, that attributable to cnc regulation, and the CNV at the locus.

**Figure 3.**
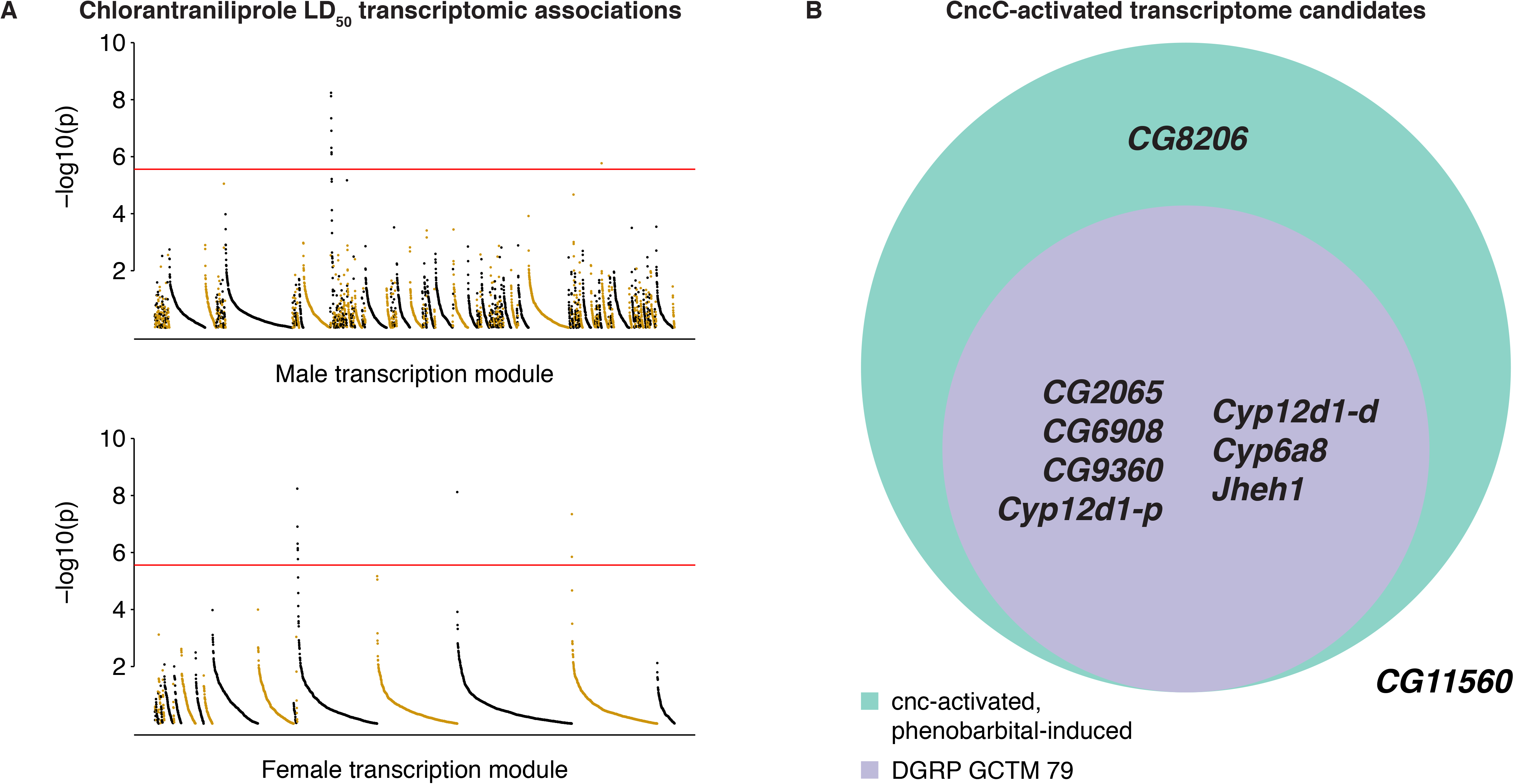
Bonferroni-significant (*p*<2.76×10^-6^) chlorantraniliprole LD_50_ phenotype to transcriptome associations. (*A*) *p*-values of association between the chlorantraniliprole LD_50_ phenotype and levels of 18139 transcripts, grouped into sex-specific genetically correlated transcriptional modules (GCTM) The transcripts within each module are ordered by rank to give ‘opera house’ plots. (*B*) Eight of the nine transcripts are affected by ectopic *cnc* expression and seven are members of DGRP male GCTM 79.

### Validation of *Strn-Mlck* and *Cyp12d1-p* in chlorantraniliprole survivorship traits

To support the involvement of *Strn-Mlck* in chlorantraniliprole survivorship, two DGRP lines (RAL-59 and RAL-399) that differ by both LD_50_ and *Strn-Mlck* variants, were crossed. Comparison of probit curves from parental lines and the F1 suggest that the more resistant allele is recessive (degree of dominance =-0.76 at LD_50_, -0.84 at LD_90_). A pair of RFLP assays showed that this haplotype was significantly enriched in chlorantraniliprole-screened F2 survivors over untreated controls (χ^2^ *p*=9.97×10^-7^), indicating that this region is indeed associated with chlorantraniliprole survivorship. The importance of *Strn-Mlck* was also tested using three separate TRiP lines crossed to the *Actin5c* driver. In two of the three crosses, flies with *Strn-Mlck* knocked down showed significantly decreased LD_50_s relative to *CyO* siblings, (Fig. 4).

**Figure 4.**
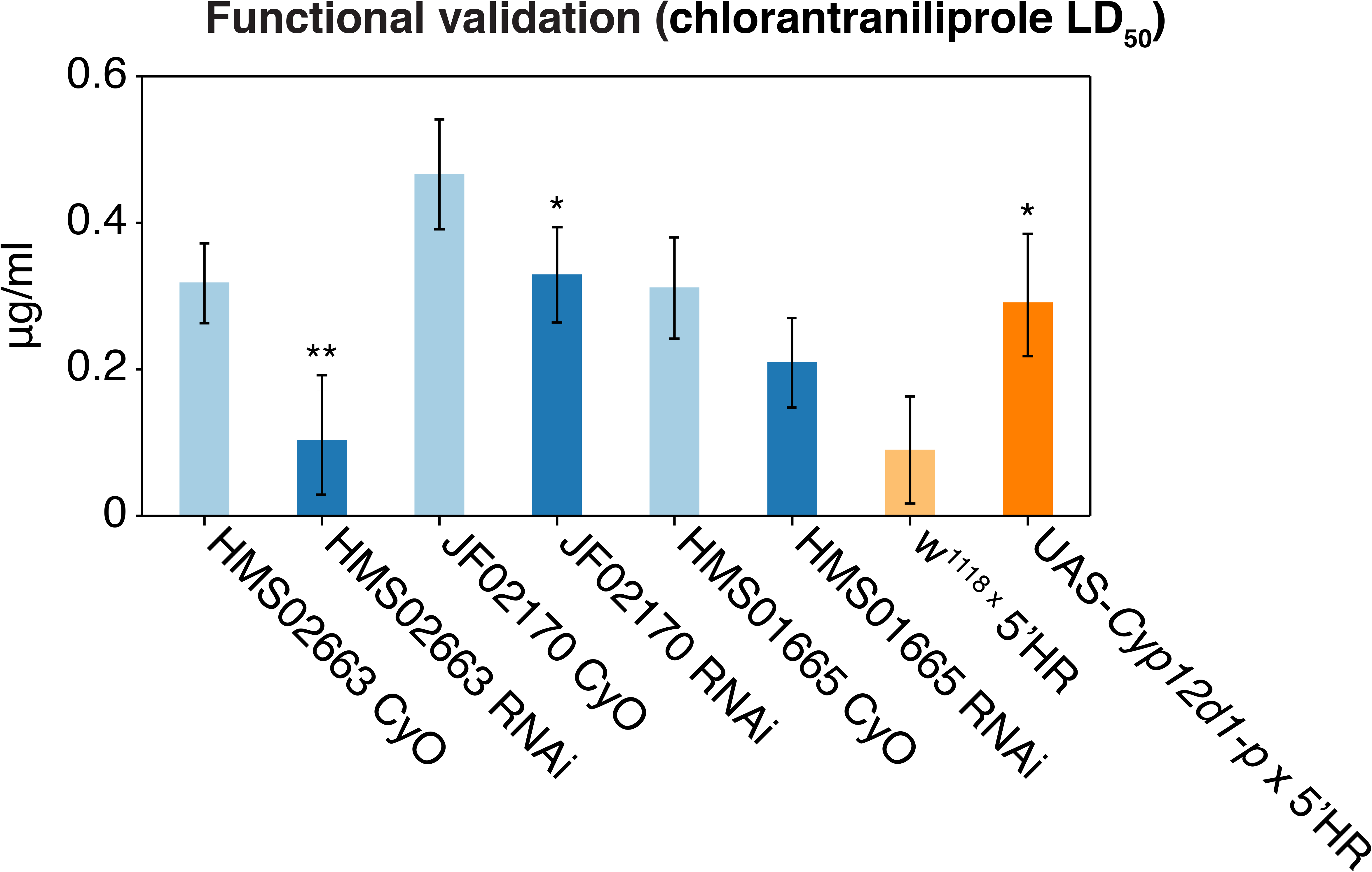
Verification of chlorantraniliprole candidates. Transgenic manipulation of candidate resistance genes. LD_50_ values of combined reciprocal crosses for RNAi knockdown of *Strn-Mlck* using three TRiP lines (HMS02663, JF02171 and HMS01665) crossed to the *Actin5c* driver. In two out of three of the crosses, *Strn-Mlck* knockdown results in a significantly decreased chlorantraniliprole LD_50_ relative to *CyO* siblings. Transgenic overexpression of *Cyp12d1-p* significantly increases chlorantraniliprole LD_50_. Error bars represent 95% confidence of probit fit at LD_50_.

The involvement of *Cyp12d1-p* was tested using the GAL4-UAS system and the *6g1*HR-GAL4 driver (36); flies overexpressing *Cyp12d1-p* in key metabolic tissues were 2.2-fold more tolerant to chlorantraniliprole than controls (Fig. 4). This system was also employed to test cross-resistance to the closely related insecticide, cyantraniliprole. At three doses, survivorship of flies overexpressing *Cyp12d1-p* was significantly increased relative to controls (*p*<0.05, two tailed t-test assuming unequal variances), suggesting that the enzyme acts on chemical moieties that the two insecticides have in common.

#### Induction of *Cyp12d1*

It has been previously demonstrated that both *Cyp12d1-p* and *Cyp12d1-d* are up-regulated in response to phenobarbital and caffeine (35). To test if chlorantraniliprole or cyantraniliprole induces *Cyp12d1* expression, *Cyp12d1* transcript levels (*Cyp12d1-p* and *Cyp12d1-d* were not distinguished) were quantified after larvae from a laboratory strain (*Canton-S*) and two DGRP lines (chlorantraniliprole-resistant RAL-59 and chlorantraniliprole-susceptible RAL-399) were exposed to chlorantraniliprole, cyantraniliprole, phenobarbital and caffeine. In accordance with DGRP transcriptome data from Huang *et al*. (30), we found *Cyp12d1* transcript abundance in unexposed larvae significantly higher in the resistant DGRP line RAL-59 relative to susceptible line RAL-399 (Fig. 5). In all three lines *Cyp12d1* transcript levels were increased significantly following exposure to phenobarbital and caffeine, but not following exposure to chlorantraniliprole or cyantraniliprole (Fig. 5).

**Figure 5.**
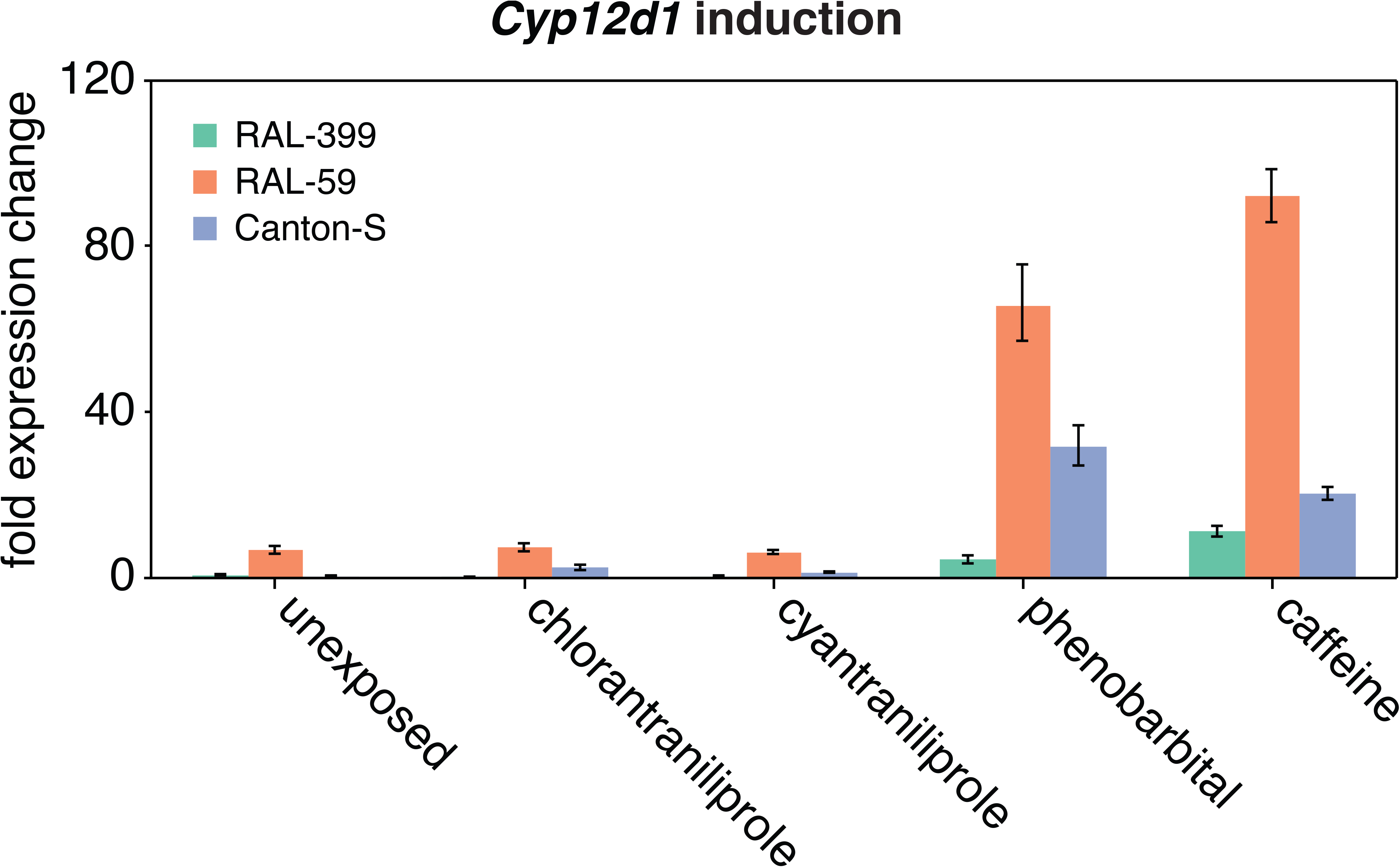
Variation and inducibility of *Cyp12d1* expression: *Cyp12d1* expression (both *Cyp12d1-p* and *Cyp12d1-d* detected; normalized to housekeeper *CG11322*) in third-instar larvae from three DGRP lines after exposure to four xenobiotic compounds. Error bars represent standard deviation of the first derivative.

## Discussion

### Genetic architecture changes with insecticide dose

Theoretical considerations have led to the prediction that the genetic architecture of insecticide resistance would become less polygenic with increasing dose of an insecticide (37). The DGRP allows identification of polygenes affecting survivorship on different doses of an insecticide in an unprecedented way. Concurring with expectations, smaller effect sizes were observed in the lowest dose GWAS. However, relatively few variants passed the significance threshold in the GWAS at this dose compared to those performed at higher doses. This is a consequence of the increasingly non-normal distribution of the survivorship among lines as dose increases and reflects the statistical mechanics and distribution assumptions of GWAS, and the degree of linkage disequilibrium around associated variants at lower frequencies. We addressed this by modelling the additive contribution of small-effect alleles using a Bayesian Linear Regression. At the three highest doses, and the dose inferred to kill 50% of flies (LD_50_), many of the associated variants are within large stretches of LD around *Strn-Mlck*. When the LD is accounted for, the number of genes associated with survivorship on high doses was indeed lower, consistent with theoretical predictions. The genetic architecture of lower dose resistance is far more polygenic, with very few of the top associations showing strong LD relationships. There is also a loss of explanatory power at 0. 5μg/ml, with R^2^ plateauing at 69%, making it obvious that the set of associated variants is not capturing all the variation in this phenotype. In addition, the heritability at the lowest dose is smaller, suggesting other environmental effects are more pronounced at this dose; or that epistasis, which is likely to be more prevalent with a greater level of polygenicity, is playing a role (38).

### *Strn-Mlck:* a novel gene of major effect

The GWAS presented here identified fifteen variants in *Strn-Mlck* gene as marking an allele of major effect. This genome-wide and unbiased approach was confirmed by two methods: RNAi-knockdown supports the potential of this gene to contribute to the trait, and linkage mapping indicates that a naturally occurring variant of major effect occurs in this vicinity. *Strn-Mlck* has one of the largest coding regions in *D. melanogaster*, spanning 38kb. A member of the Titan family, *Strn-Mlck* is highly complex, with a total of 33 exons, three start sites and three poly-A sites, leading to sixteen predicted isoforms. Myosin light chain kinases (MLCKs) are serine/threonine kinases whose substrate is the myosin regulatory chain, a component of the thick filaments in muscles. Most MLCKs are calmodulin-dependent, meaning they are proteins that are sensitive to calcium homeostasis. *Strn-Mlck’s* association with the muscle and its sensitivity to calcium, the intracellular release of which is perturbed with ryanodine insecticides, strengthen the case that *Strn-Mlck* is involved. The alleles that increase survivorship may act downstream of the insecticide binding to the ryanodine receptor, and simply ameliorate the effect of the insecticide. Alternatively, the allelic variants may act upstream by influencing the development of the nervous system such that some flies are less sensitive to the insecticide.

### Transcriptomic associations identify a known detoxification pathway

Underscoring the power of the systems approach that is afforded by the DGRP, associations between chlorantraniliprole tolerance phenotypes and gene expression levels (30) implicated a completely different set of genes from GWAS candidates. A strong association was observed between chlorantraniliprole LD_50_ and the expression of a group of genes known to be co-regulated in response to xenobiotics (35). These genes have previously been found to be regulated by cnc (35), a transcription factor well conserved across both vertebrates and invertebrates (39). cnc and its mammalian homolog Nrf2 are CNC-bZIP transcription factors that play a well-conserved role in oxidative stress response (40). While most studies characterize this pathway in terms of its induction in response to oxidative stress, the transcript-level variation identified in this study comes from unexposed flies, suggesting variation in the constitutive expression level of cnc-regulated transcripts among DGRP lines, a phenomenon that has previously been linked to insecticide resistance (35, 40).

The constitutively up-regulated cnc-regulated transcripts identified in the DGRP include potential detoxification enzymes including two cytochrome P450s (*Cyp12d1, Cyp6a8*), a juvenile hormone epoxide hydrolase (*Jheh1*), two oxidoreductases (*CG2065* and *CG9360*) and a gene containing a domain of unknown DUF227 (*CG6908*). Guio *et al*. (41) demonstrated that the insertion of a *Bari1* transposable element upstream of *Jheh1* introduces additional cnc binding motifs. This results in increased inducibility of *Jheh1* and its tandem paralog, *Jheh2* in response to oxidative stress, conferring increased resistance to the organophosphate insecticide malathion.

*Cyp12d1* induction has been associated with exposure to multiple xenobiotics, including DDT, caffeine (42), pyrethrum (43), atrazine (44), and piperonyl butoxide (PBO; 45). *Cyp12d1* is very much an expected tolerance candidate, as there is precedent for the involvement of cytochrome P450s in the metabolism of chlorantraniliprole (46), and overexpression of *Cyp12d1* has been shown to confer resistance to DDT, dicyclanil and malathion (47, 48). Due to the polymorphic status of the *Cyp12d1* in *D. melanogaster*, it has been assumed that the duplication is a recent event (49), and the two copies differentiate at three amino acids within the 3.7kb duplicated region in the *y; cn bw sp;* reference genome. There also appears to be a notable reduction of the 3’ UTR of *Cyp12d1-d*, which is speculated to affect post-transcriptional regulation (49). Duplication frequency in the DGRP is correlated with higher expression levels in adults, a pattern that is also seen in other outbred populations (50). While the DGRP transcriptome data was measured in adults, our qPCR results suggest that this difference is also observed in larvae.

As the transcription output of a gene can be affected by numerous variants, including rare variants, genes that are missed in the GWAS may be identified by phenotype to transcriptome associations. Given that the transcriptome analysis implicated *Cyp12d1* we examined its position in all the datasets more carefully. We identified two *cis*-eQTL of *Cyp12d1* (2R:6994376 and 2R:7007339; 30) associated with survivorship at 4ug/ml and 1ug/ml doses respectively. Both these variants are in LD with the duplicated state of *Cyp12d1* (R^2^=0.46 and 0.72 respectively), suggesting the effect of the *Cyp12d1* duplication was indirectly detected in the GWAS. Huang *et al*. (30) also report a *trans-eQTL* for *Cyp12d1* occurring in *sli*, the gene that ranks second in the LD_50_ GWAS after *Strn-Mlck*. Thus, trans-regulatory variation may be combining with *cis*-regulatory variation to affect the transcriptional output of *Cyp12d1*, and chlorantraniliprole survivorship.

The observation that an eQTL for a cnc-regulated gene maps to a region strongly implicated in the GWAS is intriguing. This study raises the possibility that there is a link between muscle function as implicated by the GWAS, and the oxidative stress pathway as implicated by the transcriptome associations. There is some support for this in the literature; mutations in *LamC*, a LD_50_ GWAS candidate, trigger cellular redox imbalance, an enrichment of cnc in the cytosol of larval muscle genes, and an increase in the baseline expression of genes shown to be up-regulated by ectopic *cnc* expression (25, 51).

### Conclusions

Here we set out to explore the genetic architure underlying a completely novel insecticide chemistry for which there was no expected adaptive precedence because the population sample was collected before chlorantraniliprole was deployed in the field, and is therefore completely naïve to this insecticide and indeed all of the insecticides in the new anthranilic diamide class. This contrasts to several recent studies where insecticide resistance loci have been found to feature genes of major effect that have been built through a series of adaptive substitutions (23, 52, 53). We explored the genetic architecture of chlorantraniliprole resistance at multiple doses and surprisingly found that at higher doses there was clearly a gene of major effect. Standing variation in the neuromuscular gene *Strn-Mlck* increased the LD_50_ by ~3ug/mL. This is not the first time that molecular changes underpinning insecticide resistance can be considered as exaptations (eg. 54), however this case unambiguously demonstrates that standing variation in a population is relevant to a future selective agent.

*Drosophila melanogaster* is not a pest insect but it is common in orchards where these new insecticides are being applied. It may therefore be thought of a sentinel species, such that if, in the future, there is evidence for selection at *Strn-Mlck* or the loci we identified here, then this may be attributed to the use of these new insecticides. Given the precedent for the involvement of the cnc pathway in insecticide resistance in pest species (25), perhaps the factors which cause the constitutive activation of this regulatory hub present the greatest concern.

## Materials and Methods

### Fly lines

All DGRP lines and TRiP (Transgenic RNAi Project lines; 55) were obtained from the Bloomington Drosophila Stock Centre. UAS-*Cyp12d1-p* and *6g1*HR-GAL4 were obtained from the Batterham Lab at the University of Melbourne (36). Lines were maintained on cornmeal-yeast-agar media and were kept at 25°C at constant light for at least one generation before use.

### Chlorantraniliprole phenotyping

Altacor^®^ (350 g/kg Chlorantraniliprole/Rynaxypyr™) was obtained from DuPont Australia. Wettable granules were dispersed in water to make a stock solution of 100μg/ml chlorantraniliprole. Chlorantraniliprole was administered through cornmeal-yeast-agar fly media. Insecticide was added to the media once it had cooled below 55°C. Food dye was added in conjunction with the insecticide to ensure even dispersal through the food. To set up the assay, adults from each of the DGRP lines were placed in laying cages with a 5cm diameter apple juice plate affixed to the bottom. The flies were left to lay in the cages for 24 hours. After this time, the plates were removed and replaced. Eggs were left for 24 hours to hatch and allow the development to first instar. Fifty first-instar larvae were then placed into fly vials, each containing 10ml of chlorantraniliprole-dosed media. This was done in triplicate for each line on each dose.

A total of six doses for each DGRP line were used in the initial screens (0.5, 1, 2, 3 4 and 5μg/ml), with a number of lines having to be re-screened on higher doses (6, 8 and 12μg/ml) in order to accurately determine the LD_50_. Survivorship was scored strictly eleven days after picking, with survivors being defined as any fly that successfully eclosed.

For calculation of LD_50_s for both DGRP and transgenic lines, linear models were fitted to dose-mortality data on a log-probit scale using ‘glm’ in the R statistical package (56) and scripts from Johnson *et al*. (57). 50% lethal dose (LD_50_) values and 95% confidence intervals were calculated using Fieller’s method from fitted linear models (58).

### Genome-wide Association Studies

GWAS were performed on six single-dose phenotypes and the LD_50_ for 154 DGRP lines using the DGRP pipeline (59; http://dgrp2.gnets.ncsu.edu) that corrects for the effects of *Wolbachia* infection status and five common chromosome genotypes. 1,887,900 variants were tested in each GWAS. As direct modification of the DGRP GWAS pipeline is not possible, the LD_50_ phenotype was manually corrected for the calculated effect size of associated *Strn-Mlck* variants, and resubmitted to the DGRP pipeline.

### Genomic Prediction

For genomic prediction analysis across the top variants associated with each dose phenotype and the LD_50_, a Bayesian Linear Regression coupled with LASSO was implemented using the Bayesian Linear Regression package (BLR; 60) in R, with the phenotype acting as the data vector and the genotypes the incidence matrix for βL. Up to 500 of the top variants associated with each dose phenotype (as ranked by *p*-value) were used to explain the phenotype. Each model was implemented with 5500 iterations, with a burn in of 5000 iterations and a thinning interval of 50.

### Network Based Analysis

To examine annotated interactions between different GWAS candidates, R spider software (33); http://www.bioprofiling.de/R_spider.html) was used to map GWAS candidates to KEGG and Reactome pathways.

### Phenotype to transcriptome associations

Transcriptome data for 1-3 day old adult flies from 185 DGRP lines were recovered from the DGRP website (http://dgrp.gnets.ncsu.edu/data.html; 30). Mean transcription level was calculated for each gene in each sex from two biological replicates, to give a mean level for each of the 18,140 transcripts measured by Huang *et al*. (30) in each DGRP line, for both males and females. The mean of male and female transcript levels was then calculated. A linear model was fit between mean transcription level of each gene measured by Huang *et al*. (30) and chlorantraniliprole LD_50_.

### *Strn-Mlck* RFLP mapping crosses

To examine the dominance of chlorantraniliprole resistance and to confirm the association of GWAS-associated *Strn-Mlck* variants with resistance, crosses were set up between DGRP lines RAL-399 and RAL-59. These lines have contrasting chlorantraniliprole resistance phenotypes, are free of any of the major cosmopolitan inversions (59), and carry different states at the top GWAS variants in *Strn-Mlck*. Reciprocal crosses were performed to obtain two classes of F1 progeny to check for maternal and X linked effects. Offspring from the reciprocal crosses were combined in equal numbers to establish an F2 population.

For the cross-typing, single-fly DNA extractions were performed (61). Two sets of primers for RFLPs were designed to distinguish ‘resistant’ and ‘susceptible’ *Strn-Mlck* haplotypes: EcoRV: 5’ TCAGTTCGTTGGTGTTCAGG 3’, 5’ ACTTCACGGTCAACCTGTCC 3’ and *Hpa*ll: 5’ GTTCAGATTCATCGCCATCC 3’, 5’ ACCTGCACTACCACGTACCC 3’. A total of 190 flies were scored on both assays. For each sample, 10μL PCRs were set up using 5μL of GoTaq Green (Promega, #9PIM712), 1μL each of Forward and Reverse Primers (5μL) and 3μL of sterile water.

Half of this sample was then run out on a 2% agarose gel and visualized with ethidium bromide. The remaining 5μl of PCR product was digested with either 1 μL *Eco*RV or *Hpa*ll, 1μL of CutSmart Buffer (NEB) and 3μl added sterile water. All samples were incubated at 37°C for 3 hours before being visualised on a 2% agarose gel stained with ethidium bromide.

### Knockdown of *Strn-Mlck*

RNAi knockdown of different transcripts of the *Strn-Mlck* was carried out using three different UAS lines from the TRiP Library (55; http://www.flyrnai.org), as no single line targets all of the predicted isoforms of *Strn-Mlck*. The three lines, HMS01665, HMS02663 and JF02171 target exons 1, 4 and 30 respectively. The DRSC Off-Target finder (http://www.flyrnai.org/RNAi_find_frag_free.html) predicts no additional targets for either HMS02663 or JF02171, but HMS01665 potentially targets *Cyp4aa1* upstream of *Strn-Mlck*. Crosses were carried out using the ubiquitous *actin5c* driver. Reciprocal crosses were performed for three TRiP lines. The presence of the *CyO* balancer chromosome in the *Actin5c* driver line meant that not all offspring of each cross would inherit the UAS-RNAi construct. For these crosses, the *CyO*-carrying offspring were used as controls for the UAS siblings.

### Overexpression of *Cyp12d1-p*

*Cyp12d1-p* was overexpressed using the GAL4/UAS system (62) and the *6g1*HR-GAL4 driver described by Chung *et al*. (36). *6g1*HR-GAL4 virgin females, in which GAL4 is regulated by *Cyp6g1* upstream sequence originating from Hikone-R line flies, were crossed to males carrying an additional copy of *Cyp12d1-p* under control of a UAS promoter (47). *w^1118^* was used as a control for *UAS-Cyp12d1-p*.

### *Cyp12d1* qRT-PCR induction assays

For each replicate of each line, 50 3^rd^-instar larvae were added to plates containing cornmeal-yeast-agar media and either 2.5μg/ml chlorantraniliprole, 0.5μg/ml cyantraniliprole, 10mM phenobarbital (dissolved in ETOH) or 1.5mg/ml caffeine (dissolved in 80C water). Control plates were untreated. Larvae were allowed to feed for four hours before being suspended in Trisure (Bioline) and snap frozen in liquid nitrogen for storage at -70°C. RNA was later extracted following the standard Trisure protocol. cDNA was synthesised using M-MuLV reverse transcriptase (NEB) and random nonamer primers following the standard NEB reverse transcriptase protocol. qRT-PCR was performed on the Roche Light Cycler 480 using the following primers: *Cyp12d1* (*Cyp12d1-p* or *Cyp12d1-d*): 5’ GGGGAAAACTACGATCAGCC 3’, 5’ CGGATTTCCTTAATGCGCTCT 3’, *CG11322* (housekeeper): 5’ TGGGCAGTGCCTTCTACATTT 3’, 5’ CGTACGCACCTCGCTTGTT 3’.

## Acknowledgements

We would like to acknowledge Trudy Mackay for advice and wisdom, Stephen Wilcox for his assistance with molecular biology analyses, Phil Batterham for UAS lines and discussion, Josefa González for discussion and John McKenzie for comments on the manuscript. Some of this work was supported by funding from the Australia Research Council-DP0985013.

## Figure Legends

**Figure S1**. The explanatory power of phenotype-associated DGRP variants for each chlorantraniliprole dose contrasts to the number of associated variants. For each dose the number of variants that cross the GWAS significance threshold are shown as black dots (y-axis on the right). The bar graphs report the explanatory power (y-axis on left) for 5, 10, 25 and 50 variants for each dose using a genomic prediction. This shows that more genes are required to explain the genetic architecture of the lowest dose. For example, if the top 50 most-associated variants of the 0.5ug/mL dose are considered they explain about the same amount of the phenotypic variation (R=0.43) as the top 5 variants of the 5ug/mL dose.

**Figure S2**. q-q plot showing that correction of the LD_50_ phenotype for the effect of *Strn-Mlck* does not return the observed GWAS *p*-values to expected levels.

**Figure S3**. R spider networks. R spider networks of GWAS candidates (*p*<1×10^-5^) from LD_50_ (A) and *Strn-Mlck*-corrected LD_50_ phenotypes (*B*). GWAS candidates are annotated with minimum *p*-value (black), maximum MAF (red) and maximum effect size (blue).

