## Supplementary figures and images for "*Cis* and *trans*-acting variants contribute to survivorship in a naïve *Drosophila melanogaster* population exposed to ryanoid insecticides"

### Supp fig 1

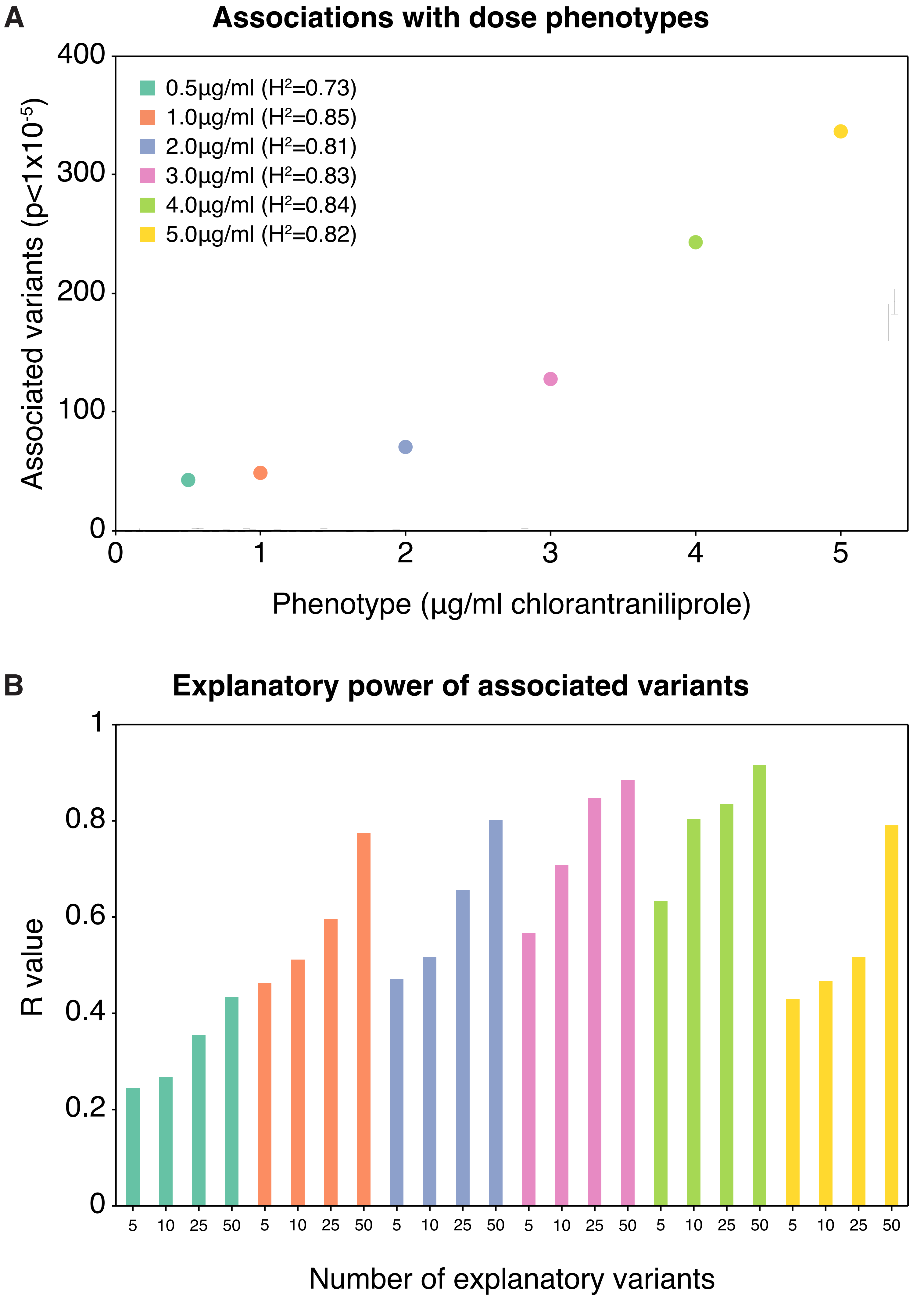

### Supp fig 2

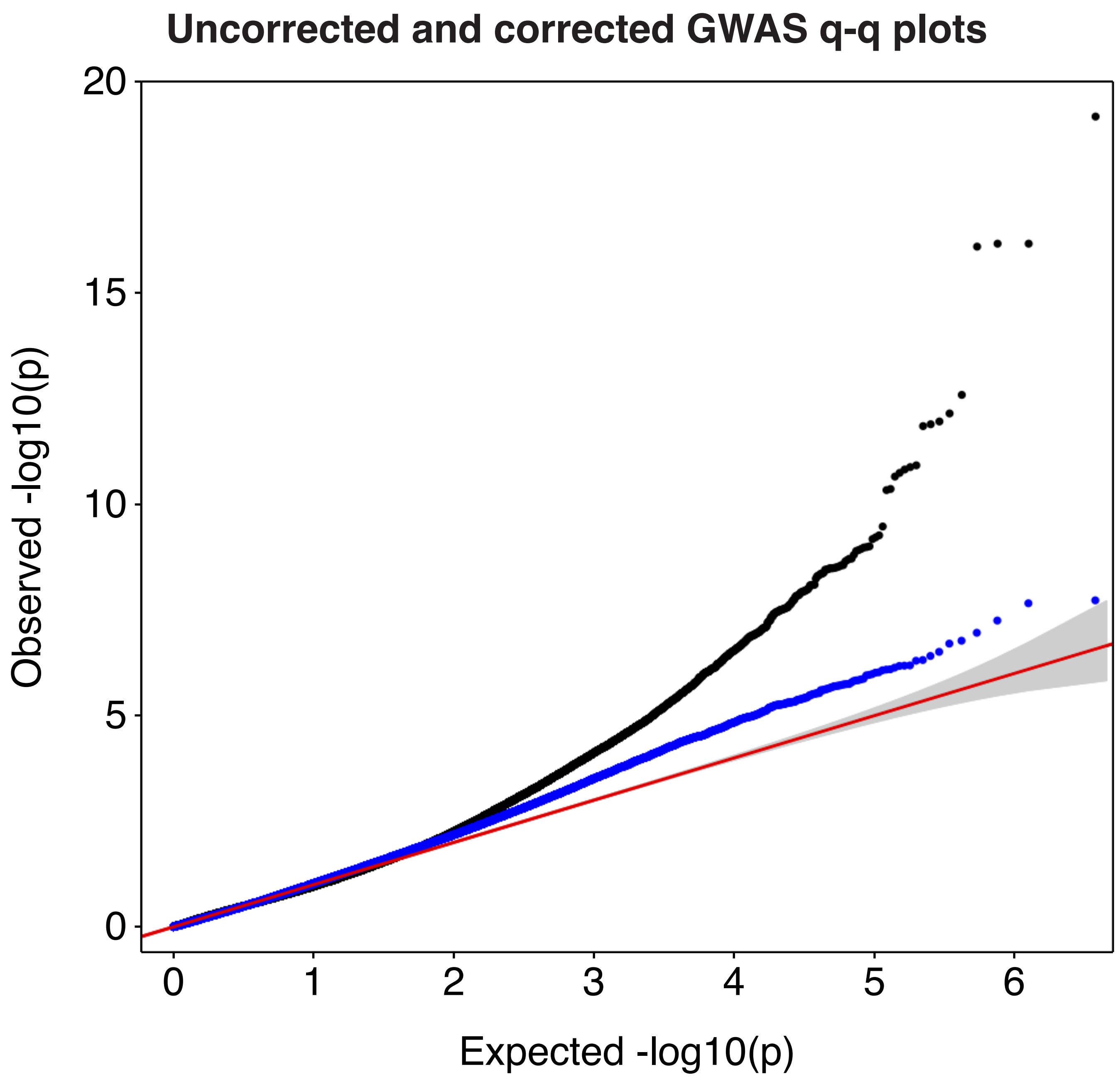

### Supp fig 3

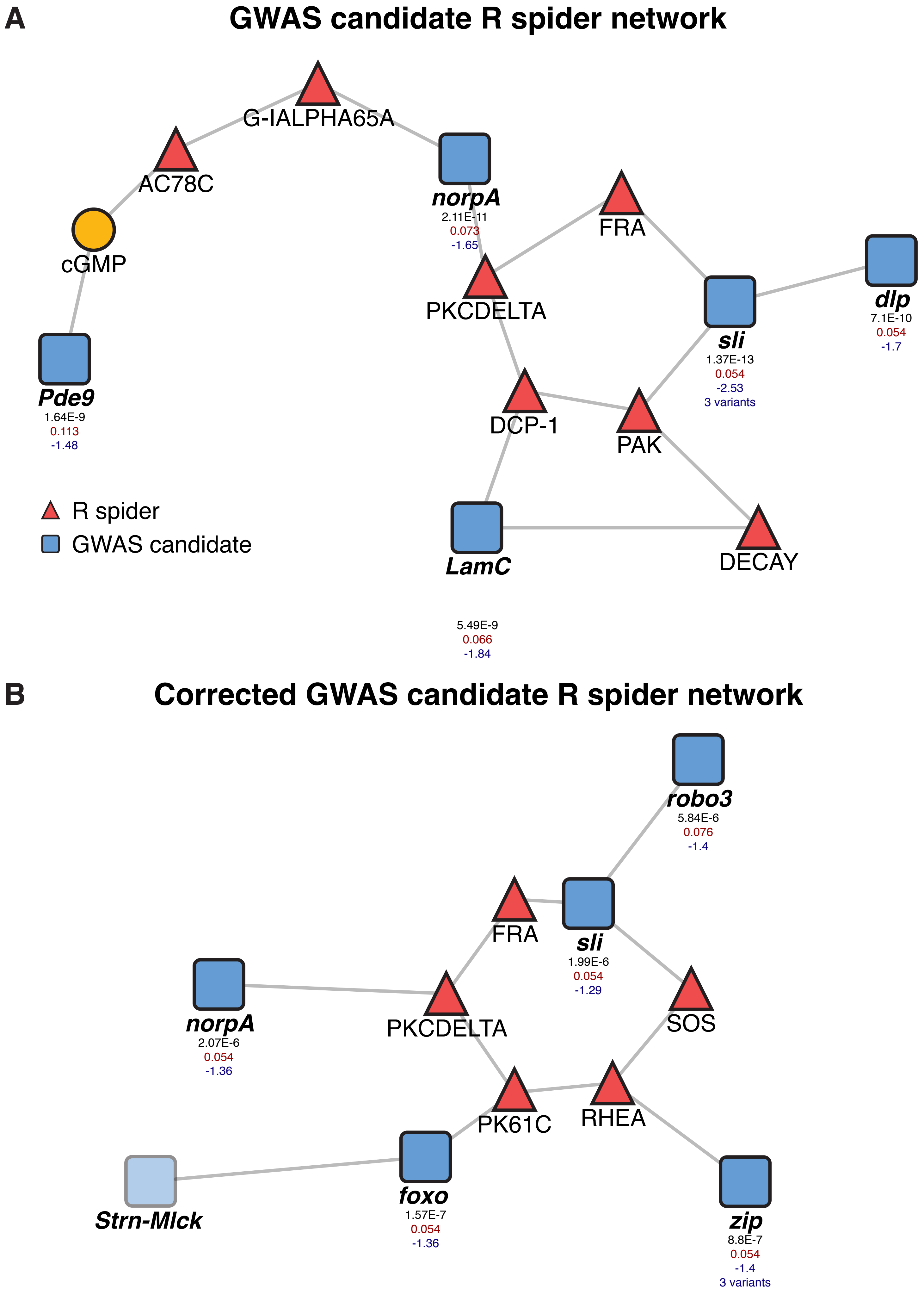
